# Zebrafish DANA retroposon can form large zebrafish sequence in human Hepg2 and 293T cell lines

**DOI:** 10.1101/493155

**Authors:** Zheming Cao, Weidong Ding, Jun Qiang, Xuwen Bing, Pao Xu

## Abstract

In this study, we cloned small zebrafish retroposon DANA from zebrafish genome and constructed the lentiviral expression vector pEB-GFP (T2A)PURO. Three human cell lines including 293T, Hepg2 and LO2 were selected as infection targets. After detecting the expression of DANA, we found that the expression of DANA retroposon in three cells had different effects on cell lines through chromosome walking. Among these cells, LO2 showed no DANA retrotrans-position, while 293T and Hepg2 cell lines displayed retrotrans-position with the formation of some zebrafish genome fragments. Thereafter, we constructed a mutant of DANA retroposon and infected it in 293T cells, but no retrotrans-position was found after chromosome walking. Re-sequencing of the two cell lines (293T and Hepg2) showed that a large number of zebrafish genome fragments were found in the genomes of both cell lines, which could be divided into four types. The first type was zebrafish microsatellite sequence, accounting for 79.23% and 74.45% in 293T cell line and Hepg2 cell line, respectively. The second type was the sequence with a small amount of poly A or T, and the third type was the sequence with poly G or C, and the second and third types accounted very low proportion. The fourth type was composed of coding sequence and non-coding sequence, with large difference and very low proportion of common sequences between the two cell lines. Taken together, this study indicated that zebrafish DANA retroposon can result in retrotrans-position using the retrotrans system of human cell lines.

## 1. Introduction

There are large numbers of retroposon sequences in the genome of eukaryotic higher organisms. For example, only 3% of the sequences of human genome are exon sequences, while the transferable components account for more than 35% of the total sequences of human genome (1, 2). Izsvak Z et al. (3) found that DANA retroposon is a type of short interspersed repetitive element (SINE), which is from zebrafish genome. DANA related sequence accounted for nearly 10% of zebrafish genome, however, its distribution has not been found in other carp fishes (3).

SINE sequence (SINES) belongs to the nonviral superfamily of retroposon, and it is the transcription of RNA polymerase III transcription, widely distributing in the genome of many eukaryotic higher organisms. The largest SINES in the human genome is the Alu family, with approximately 300,000 members (4, 5). Related sequences also exist in mice with 50,000 members known as B1 families, Chinese rats with Alu equivalent families and other animals. Some members of the Alu family can be transcribed into RNA (6, 7).

The expression of ALu sequence can be detected in human cells, and it can also lead to retrotrans-position. Moreover, SINES is the transcription of RNA polymerase III, and it can result in retrotrans-position using the ability of other retroposon in cells. Thus, we assumed that whether retrotrans-position could exist if we express zebrafish DANA retroposon in human cell lines.

## 2. Materials and methods

### 2.1. The acquisition of DANA retroposon

Common wild type zebrafish was used in this study. The genome was extracted using the genome extraction kit (Takara) according to the instructions. The primers used for DANA amplification were GGCGACRCAGTGGCGCAGTRGG and TTTTCTTTTTGGCTTAGTCCC. The PCR procedure was 94 °C for 2 min, 25 cycles of 94 °C for 30 s, 55 °C for 45 s and 72 °C for 1 min, and 72 °C for 2 min.

### 2.2. The design of DANA retroposon mutant

Based on the results of Z Izsvak et al. (3), the DANA retroposon mutant was designed by modifying two Poly III liked promoter sequences in the conserved sequence 1 and 4 and a fragment in the conserved sequence 2.

### 2.3. Virus packaging

DANA gene was constructed into the lentiviral expression vector to form CMV-MCS-EF1-copGFP-T2A-Puro vector and the vector was then used for lentivirus packaging by Guangzhou Huijun Biotechnology Co. LTD.

### 2.4. Cell culture

After removing the medium, 1 ml of 0.25% EDTA-trypsin solution was added to cover the whole bottom of the culture bottle and kept for 5 min with dynamic monitoring by the microscope. Thereafter, 2 fold volumes of DMEM medium were added to terminate the digestion, to blow and to suspend the cells. After centrifugation at 1000 rpm for 3min, the supernatant was discarded and the culture medium was added to blow the precipitate and form the resuspended cells. Finally, appropriate cells were inoculated into a new culture bottle and cultured in the cell incubator with 5% of CO_2_.

### 2.5. Lentivirus transfecting into adherent cell

At 18 to 24 hours before the transfection, adherent cells were put into the 6 orifice plate with 5 × 10^5^ cells in each hole to let the cell number to 1× 10^6^ in each hole in the lentivirus transfection. The next day, 2 ml fresh medium containing 6 μg/ml polybrene was added to replace the original medium, and an appropriate amount of virus suspension was added to incubate at 37 °C for 4 hours. Thereater, 2 ml fresh medium was added to dilute the polybrene. After culture for 24 hours, fresh medium was added to replace the culture medium containing virus for further culture.

### 2.6. Collection of cell sample and DNA extraction

After removing the medium, 1 ml of 0.25% EDTA-trypsin solution was added to cover the whole bottom of the culture bottle and kept for 5 min with dynamic monitoring by the microscope. Thereafter, 2 fold volumes of DMEM medium were added to terminate the digestion, to blow and to suspend the cells. After centrifugation at 1000 rpm for 3min, the supernatant was discarded and PBS solution was added to blow the precipitate and form the resuspended cells. Then the resuspended cells were used for DNA extraction and Trizol treatment. After centrifugation at 1000 rpm for 3min, the supernatant was discarded and 500 μl of Trizol was added in the EP tube. After quick freezing by liquid nitrogen, the samples were stored in the refrigerator at −80 °C.

### 2.7. RNA extraction and cDNA synthesis

RNA extraction and further cDNA synthesis were performed using RNAiso Plus (Code No.9108Q, Takara) and PrimeScript ™ II 1st Strand cDNA Synthesis Kit (Code No.6210A, Takara), respectively, according to the instructions.

### 2.8. Semi-quantitative PCR analysis

Human β-actin (cctagaagcatttgcggtgg; gagctacgagctgcctgacg) was selected as the internal reference gene. PCR reaction system was composed with 2 μl of cDNA, 2 μl of 10 pM forward primer, 2 μl of 10 pM reverse primer, 4 μl of 2 mM dNTP, 5 μl of 10×PCR buffer, 1 μl of Taq enzyme, and 34 μl of double distilled water. The PCR procedure was 94 °C for 2 min, n cycles of 94 °C for 30 s, 55 °C for 45 s and 72 °C for 1 min, and 72 °C for 2 min. Then the PCR product was examined by agarose gel electrophoresis and was visualized under ultraviolet lamp.

### 2.9. Chromosome walking analysis

Chromosome walking was performed using Genome Walking Kit (Code No.6108, Takara), according to the instructions. The amplified product was recovered using the Gel Recovery Kit (Shennengbocai), cloned and sequenced by Baoshang Biological Company.

### 2.10. Re-sequencing of the human genome

The genome of human cell lines was re-sequenced by Shenzhen Huada Ocean Technology Co., LTD. The reference genome version was Danio_rerio.GRCz10.dna.toplevel.fa, which was downloaded from ftp://ftp.ensembl.org/pub/release-91/fasta/danio_rerio/dna/Danio_rerio.GRCz10.dna.toplevel.fa.

## 3. Results

### 3.1. DANA sequence

The DANA sequence was amplified from the common zebrafish genome. After DNA clone and sequencing, a series of DANA sequences with slightly difference were obtained. Thereafter, the following sequence was chosen for expression. TTTTTTCTTTTTGGCTTAGTCCCTTTATTATTCTGGGGTCGCCACAGCCCAATGAACCACCAACTTTTCCAGCATATCTTTTACGCAGCGGATGCCCTTCCAGCTGCAACCCATCAATGGGAAATGCCAATACACTCCCATTCACACACATACACTACGAACAATTTAGCCTACCCAATTCACTTGTACCGCATGTCTTTGGACTGTGGGGGAAACCAGAGCACCCGGAGGAAACCCATGCGAACACGGCGAGAACATGCAAACTCCACACAGAAATGCCAACTGACCCAGCCGAGACTCGAACCAGCGATCTTCTTGCTGTGAGGCAATCGTGCTACCTACTGCGCCACTGT GTCGCC

### 3.2. The analysis of cell images after virus infection

**Figure 1.**
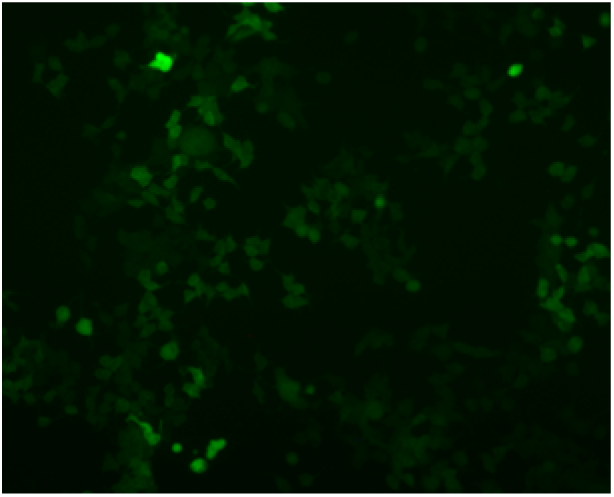
The image of virus infection in 293T cells (20th generation).

**Figure 2.**
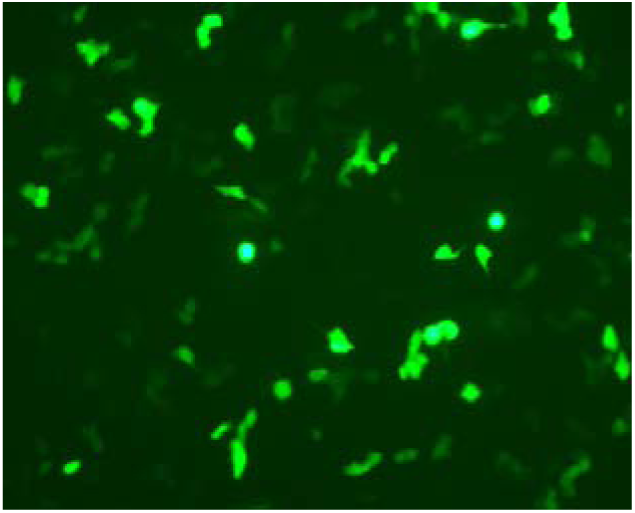
The image of virus infection in Hepg2 cells (20th generation).

**Figure 3.**
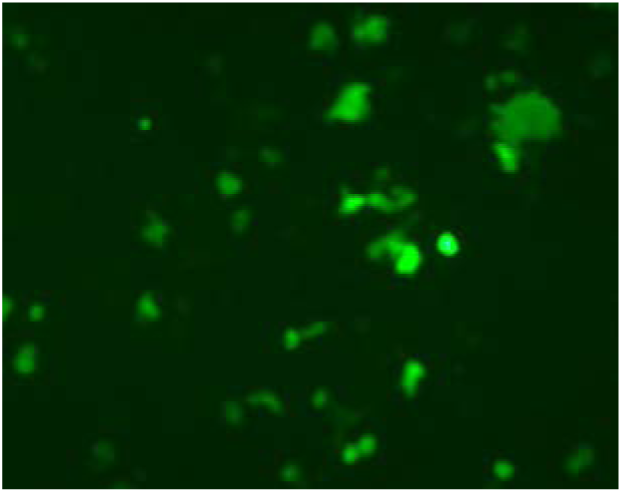
The image of virus infection in LO2 cells (20th generation).

### 3.3. Semi-quantitative PCR analysis

After choosing the best cycle number and adjusting the concentration of PCR template using *actin* gene as internal inference gene, the expression of DANA in different generations were analyzed by semi-quantitative PCR. The expression of DANA in three cell lines with the cycle of 1, 5, 10, 15, and 20 showed that no significant difference was exhibited in different generations and in different cell lines.

### 3.4. Chromosome walking analysis

Genome walking analysis in upstream and downstream of DANA retroposon showed that the upstream and downstream of LO2 cells were the sequences of packaged plasmids, indicating that the expression of DANA in this cell line had no effect on the cells and no retrotrans-position was exist. In 293T and Hepg2 cell lines, we found that there were a large number of zebrafish sequences in the upstream and downstream of DANA retroposon. The zebrafish sequences formed in 293T cell line included DKEY-259p11 in linkage 8 (117608-118551), DKEY-153N92 in linkage 2 (177824-178118), DKEY-7F16 in linkage 3(4571-3719), DKEY-238m4 in linkage 16(29645-30316), DKEY-7F16 in linkage 3(3053-3977), and DKEY-35I13 in linkage 15(112086-112469). The zebrafish sequences formed in Hepg2 cell line included DKEY-259p11 in linkage 8 (117608-118551,118860-117911,118389-118860), CH73-317m8 in linkage 20(86054-86497, 88179-87637,87432-87187). Among these sequences, DKEY-259p11 in linkage 8 (117608-118551) was commonly distributed in the two cell lines.

### 3.5. Analysis of DANA retroposon mutant

TTTTTTCTTTTTaatccgactttTTTATTATTCTGGGGTCGCCACAGCCCAATGAACCACCAACTTTTCCAGCATATCTTTTACGCAGCGGATGCCCTTCCAGCTGCAACCCATCAATGGGAAATGCCAATACACTCCCATTCACACACATACACTACGAACAATTTAGCCTACCCAATTCACTTGTACCGCATGTCTTTGGACTGTaaaaagggttAGAGCACCCGGAGGAAACCCATGCGAACACGGCGAGAACATGCAAACTCCACACAGAAATGCCAACTGACCCAGCCGAGACattcggatctGATCTTCTTGCTGTGAGGCAATCGTGCTACCTACTGCGCCACTGTGTCGCC

In order to verify whether the above results were correct, we designed the DANA retroposon mutant, and the red part is the mutated region. After lentivirus packaging, infection in 293T cell line and cultivation for 20 generations, genomic walking was performed in the upstream and downstream of the mutant. It was found that the upstream and downstream of 293T cell line were the sequences of packaged plasmids, indicating that the expression of mutant in this cell line had no effect on the cells without retrotrans-position.

### 3.6. Re-sequencing of human genome

In order to further investigate the effect of DANA retroposon on 293T and Hepg2 cells lines in zebrafish, the whole genome of two cell lines cultured for 20 generations were re-sequenced and compared the sequence of zebrafish genome. The statistical results were shown in Table 1. In the re-sequencing results of 293T cell line, a total of 1512 sequences were compared in the zebrafish genome, including 1,198 microsatellite sequences, 99 poly A or T sequences, 9 poly G or C sequences, and 206 other sequences. In Hepg2 cell line, a total of 1491 sequences were compared in zebrafish genome, including 1110 microsatellite sequences, 88 poly A or T sequences, 7 poly G or C sequences, and 286 other sequences.

The other sequences were composed of coding sequences and non-coding sequences. The statistics of coding sequences were shown in Table 2. Generally, 7 coding sequences were found in 293T cells and 22 coding sequences were found in Hepg2 cells. Only the ribosomal RNA encoded on chromosome 5 was common in both cell lines, and the others were completely different. Some coding sequences in Hepg2 cells could identified highly similar coding sequences in human, including nuclear factor I C (NFIC), SKI proto-oncogene, myosin light chain kinase, DEAD-box helicase 3, X-linked (DDX3X), kelch like family member 28, DLG associated protein 2, and piccolo presynaptic cytomatrix protein. Two coding sequences in 293T cells including nebulin (NEB) and myosin heavy chain 1could identified highly similar coding sequences in human. The others failed to find the comparison results or found the coding sequences with low similarity.

Non-coding sequences were also divided into two types, the first type with more than 5 copies distribution in the zebrafish genome (Table 3), and the second type with less than five copies. Some fragments had no significant comparison results in human genome, but had corresponding fragments in carp genome. The first type sequences were shown in Table 3, 22 and 20 sequences with more than 5 copies distribution were found in 293T cells and Hepg2 cells, respectively. After comparing these fragments with the human genome, we found that most of them were spliced together from sequences with very low similarity.

## 4. Discussion

Initially, the constructed lentivirus particles of zebrafish DANA retroposon were microinjected into the carp zygoteink, and the expression of this retroposon and the unusual retrotrans-position phenomenon were detected in the two lived carps. However, the offspring of the two carp could not survive. In order to further verify the ability of the retroposon, human 293T cells were chosen for the experiment, and it was found that the retroposon resulted in unusual retrotrans-position phenomenon in 293T cells. In order to avoid the error by pollution, this study was performed in College of Medicine in Tongji University.

This study indicated that DANA retroposon from zebrafish still had the ability of retrotrans-position in human cell lines. Due to the difference between DANA retroposon and human Alu sequence, it is still elusive whether the retrotrans-position was resulted form existing human LINE.

There have been some speculations about the mechanism of retrotrans-position. However, the result in this study is completely inconsistent with previous speculations on the mechanism of retrotrans-position. Because different cell lines could form various sequences of retroposon, so it was difficult to find obvious rules. However, some of the sequences were obviously fragments that transcribed from modified human mRNAs. We suspected that the retroposon might combine with the cellular endogenous reverse transcriptase after transcribing into RNA and interact with other RNAs or even a lot of RNA degradated pieces at the same time, and further influenced the pairing mechanism of reverse transcriptase, resulting in the human RNAs reverse transcribed into other DNA sequences and further enter into the genome during the process of reverse transcription. Zebrafish ancestors probably did not have these sequences before obtaining the retroposon. After obtaining the retroposon, many such fragments were formed in the genome by the same mechainsm. That is to say, the zebrafish sequence examined in this study might not come from the ancestral genome of zebrafish, but was caused by the unusual retrotrans-position.

This study showed that expression DANA retroposon formed many fragments in human genome that could be found in zebrafish genome, and multiple fragments were widely distributed in zebrafish genome. Then we tend to think that DANA retroposon might play an extremely important role in the formation of zebrafish specie. Because DANA retroposon could express in human cell lines and display similar function, once the retroposon was introduced into human genome, human genome might be reshaped. The expression of DANA retroposon could form zebrafish fragments in human genome and modify many coding sequences, indicating that the retroposon might involve in the expression regulation of many genes and even the modification of gene sequences.

Moreover, this study found that the retroposon resulted in no retrotrans-position in human normal liver cell line LO2, but a large number of zebrafish genome sequences in 293T and Hepg2 cell lines. This result suggested that there was no condition for the retrotrans-position in LO2 cells, while 293T and Hepg2 had such condition. Whether it was due to the non-expression of reverse transcriptase in LO2 cells or other reasons needs to be further investigated. If the phenomenon could be confirmed in other cells, it might provide a new cancer therapy.

