## Supplementary material for "Zebrafish DANA retroposon can form large zebrafish sequence in human Hepg2 and 293T cell lines": Table 1

Table 1 Comparison of resequencing results of two cell lines

| Chromosome |  | 293 | Hepg | Chromosome |  | 293 | Hepg |
| --- | --- | --- | --- | --- | --- | --- | --- |
| 1 | Microsatellite sequence | 59 | 37 | 2 | Microsatellite sequence | 53 | 40 |
|  | Poly A or T sequences | 6 | 8 |  | Poly A or T sequences | 3 | 2 |
|  | Poly G or C sequences | 1 | 1 |  | Poly G or C sequences | 0 | 0 |
|  | Other sequences | 9 | 16 |  | Other sequences | 2 | 10 |
|  | Total number of sequences | 75 | 62 |  | Total number of sequences | 58 | 52 |
| 3 | Microsatellite sequence | 46 | 48 | 4 | Microsatellite sequence | 27 | 34 |
|  | Poly A or T sequences | 5 | 2 |  | Poly A or T sequences | 3 | 2 |
|  | Poly G or C sequences | 0 | 0 |  | Poly G or C sequences | 0 | 0 |
|  | Other sequences | 7 | 14 |  | Other sequences | 6 | 3 |
|  | Total number of sequences | 58 | 64 |  | Total number of sequences | 36 | 39 |
| 5 | Microsatellite sequence | 71 | 77 | 6 | Microsatellite sequence | 67 | 60 |
|  | Poly A or T sequences | 2 | 1 |  | Poly A or T sequences | 4 | 4 |
|  | Poly G or C sequences | 0 | 0 |  | Poly G or C sequences | 0 | 0 |
|  | Other sequences | 7 | 13 |  | Other sequences | 6 | 15 |
|  | Total number of sequences | 80 | 91 |  | Total number of sequences | 77 | 79 |
| 7 | Microsatellite sequence | 77 | 60 | 8 | Microsatellite sequence | 46 | 38 |
|  | Poly A or T sequences | 14 | 9 |  | Poly A or T sequences | 1 | 0 |
|  | Poly G or C sequences | 0 | 0 |  | Poly G or C sequences | 1 | 1 |
|  | Other sequences | 13 | 16 |  | Other sequences | 11 | 13 |
|  | Total number of sequences | 104 | 85 |  | Total number of sequences | 59 | 52 |
| 9 | Microsatellite sequence | 51 | 53 | 10 | Microsatellite sequence | 38 | 34 |
|  | Poly A or T sequences | 6 | 5 |  | Poly A or T sequences | 3 | 0 |
|  | Poly G or C sequences | 2 | 1 |  | Poly G or C sequences | 0 | 0 |
|  | Other sequences | 8 | 20 |  | Other sequences | 9 | 8 |
|  | Total number of sequences | 67 | 79 |  | Total number of sequences | 50 | 42 |
| 11 | Microsatellite sequence | 37 | 43 | 12 | Microsatellite sequence | 43 | 45 |
|  | Poly A or T sequences | 6 | 4 |  | Poly A or T sequences | 5 | 3 |
|  | Poly G or C sequences | 2 | 2 |  | Poly G or C sequences | 1 | 0 |
|  | Other sequences | 8 | 9 |  | Other sequences | 6 | 11 |
|  | Total number of sequences | 53 | 58 |  | Total number of sequences | 55 | 59 |
| 13 | Microsatellite sequence | 59 | 64 | 14 | Microsatellite sequence | 42 | 36 |
|  | Poly A or T sequences | 6 | 1 |  | Poly A or T sequences | 5 | 6 |
|  | Poly G or C sequences | 0 | 0 |  | Poly G or C sequences | 0 | 0 |
|  | Other sequences | 7 | 16 |  | Other sequences | 4 | 13 |
|  | Total number of sequences | 72 | 81 |  | Total number of sequences | 51 | 55 |
| 15 | Microsatellite sequence | 53 | 45 | 16 | Microsatellite sequence | 47 | 45 |
|  | Poly A or T sequences | 2 | 1 |  | Poly A or T sequences | 6 | 4 |
|  | Poly G or C sequences | 1 | 1 |  | Poly G or C sequences | 0 | 0 |
|  | Other sequences | 9 | 13 |  | Other sequences | 8 | 14 |
|  | Total number of sequences | 65 | 60 |  | Total number of sequences | 61 | 63 |
| 17 | Microsatellite sequence | 48 | 44 | 18 | Microsatellite sequence | 54 | 63 |
|  | Poly A or T sequences | 9 | 6 |  | Poly A or T sequences | 6 | 7 |
|  | Poly G or C sequences | 0 | 0 |  | Poly G or C sequences | 0 | 0 |
|  | Other sequences | 9 | 12 |  | Other sequences | 7 | 15 |
|  | Total number of sequences | 66 | 62 |  | Total number of sequences | 67 | 85 |
| 19 | Microsatellite sequence | 48 | 41 | 20 | Microsatellite sequence | 61 | 44 |
|  | Poly A or T sequences | 5 | 5 |  | Poly A or T sequences | 2 | 1 |
|  | Poly G or C sequences | 0 | 0 |  | Poly G or C sequences | 0 | 0 |
|  | Other sequences | 9 | 8 |  | Other sequences | 7 | 12 |
|  | Total number of sequences | 62 | 54 |  | Total number of sequences | 70 | 57 |
| 21 | Microsatellite sequence | 33 | 33 | 22 | Microsatellite sequence | 23 | 24 |
|  | Poly A or T sequences | 3 | 4 |  | Poly A or T sequences | 5 | 4 |
|  | Poly G or C sequences | 0 | 0 |  | Poly G or C sequences | 0 | 0 |
|  | Other sequences | 6 | 2 |  | Other sequences | 7 | 7 |
|  | Total number of sequences | 42 | 39 |  | Total number of sequences | 35 | 35 |
| 23 | Microsatellite sequence | 38 | 36 | 24 | Microsatellite sequence | 51 | 42 |
|  | Poly A or T sequences | 4 | 2 |  | Poly A or T sequences | 5 | 5 |
|  | Poly G or C sequences | 1 | 1 |  | Poly G or C sequences | 0 | 0 |
|  | Other sequences | 5 | 7 |  | Other sequences | 10 | 10 |
|  | Total number of sequences | 49 | 46 |  | Total number of sequences | 66 | 57 |
| 25 | Microsatellite sequence | 26 | 24 |  |  |  |  |
|  | Poly A or T sequences | 4 | 2 |  |  |  |  |
|  | Poly G or C sequences | 0 | 0 |  |  |  |  |
|  | Other sequences | 3 | 8 |  |  |  |  |
|  | Total number of sequences | 33 | 34 |  |  |  |  |
