## Supplementary material for "Zebrafish DANA retroposon can form large zebrafish sequence in human Hepg2 and 293T cell lines": table 2

Table 2. Fragments with coding or transcriptional functions in zebrafish genome

| Chromosome | 293 | Hepg2 |
| --- | --- | --- |
| 1 | 0 | 0 |
| 2 | 0 | 0 |
| 3 |  | nuclear factor I/Xb (nfixb), transcript variant X12 |
| 4 | 0 | 0 |
| 5 | zgc:158463, ribosomal RNA | zgc:158463, ribosomal RNA  si:ch211-106a19.1 , transcript variant X2, misc_RNA  neuronal tyrosine-phosphorylated phosphoinositide-3-kinase adapter 1, transcript variant X1 |
| 6 | 0 | serine/arginine-rich splicing factor 3a (srsf3a), transcript variant X3, misc_RNA |
| 7 | 0 | 0 |
| 8 | 0 | v-ski avian sarcoma viral oncogene homolog a (skia), mRNA |
| 9 | nebulin (neb), transcript variant X23, mRNA  sorting nexin 4 (snx4), mRNA | myosin, light chain kinase a (mylka), transcript variant X3, mRNA  speckle-type POZ protein-like a (spopla), mRNA  interleukin 1 receptor accessory protein-like 1a (il1rapl1a), transcript variant X12, mRNA  fibroblast growth factor 14 (fgf14), transcript variant X4, mRNA  DEAD (Asp-Glu-Ala-Asp) box helicase 3a (ddx3a), transcript variant X15, mRNA  POU class 2 homeobox 1b (pou2f1b), transcript variant X9, mRNA  low density lipoprotein receptor-related protein 1Bb (lrp1bb), transcript variant X4, mRNA |
| 10 | 0 | 0 |
| 11 | 0 | 0 |
| 12 |  | si:dkey-7e14.3 , mRNA |
| 13 |  | uncharacterized LOC110440175 , ncRNA |
| 14 | 0 | 0 |
| 15 | 0 | nuclear receptor interacting protein 1a (nrip1a), transcript variant X1 |
| 16 |  | dual-specificity tyrosine-(Y)-phosphorylation regulated kinase 1B (dyrk1b), transcript variant X4 |
| 17 | 0 | kelch like family member 28 (klhl28)  somatostatin receptor 1a (sstr1a)  discs, large (Drosophila) homolog-associated protein 2a (dlgap2a), transcript variant X3 |
| 18 | 0 | piccolo presynaptic cytomatrix protein b (pclob), transcript variant X6 |
| 19 | 0 | bloodthirsty-related gene family, member 24 (btr24)  si:ch211-233f16.1 , transcript variant X4 |
| 20 | eph receptor A7 (epha7), transcript variant X1 | 0 |
| 21 | crumbs family member 2a (crb2a), mRNA | 0 |
| 22 | myosin heavy chain, fast skeletal muscle-like , misc_RNA | 0 |
| 23 | 0 | 0 |
| 24 | component of oligomeric golgi complex 7 | 0 |
| 25 | 0 | 0 |

Chromosome3：

Hepg

nuclear factor I/Xb (nfixb), transcript variant X12

AAGGGCAAGATCCGCCGCATCGACTGCCTGCGGCAAGCCGACAAGGTCTGGCGCCTGGACCTGGTGATGGTTATCCTGTTCAAGGGAAGCCCGCTGGAGAGCACGGATGGAGAGAGGCTGGTCAAGTCCCCGCAGTGCTCAAACCACGGCCTCTGCGTGCAGCCGCACCACATCGGAGTGTCCGTCAAGGAACTGGACCTGTATCTGGCCTACTTCGTCCACACGCCAG

Homo sapiens nuclear factor I C (NFIC), transcript variant X5, mRNA

Chromosome5：

293

zgc:158463 (zgc:158463), ribosomal RNA

AACGGGTTACCCGCGCCTCTCGGCGCAGGGTAGGCACACGTTGATCCGCCCATTGTGGCGCGCGTGCAGCCCCGGACATCTAAGGGCATCACAGACCTGTTATTGCTCCATCTCGCGTGGCTGAACGCCACTTGTCCCTCTAAGAAGTTGGGACGCCGACCGCGCGGGGCCGCGTAACTATTTAGCATGCCGGAGTCTCGTTCGTTATCGGAATGAACCAGACAAATCGCTCCACCAACTAAGAACGGCCATGCACCACCACCCACAGAATCGAGAAAGAGCTATCAATCTGTCAATCCTTTCCGTGTCCGGGCCGGGTGAGGTTTCCCGTGTTGAGTCAAATTAAGCCGCAGGCTCCACTCCTGGTGGTG

Hepg

zgc:158463 (zgc:158463), ribosomal RNA

CGGCGCTCCGCCAGGGCCCGGCGAGGAGCCCCGGCGGGGCCGATCCGAGGACCTCACTAAACCATCCAATCGGTAGTAGCGACGGGCGGTGTGTACAAAGGGCAGGGACTTAATCAACGCAAGCTTATGACCTGCGCTTACTGGGAATTCCTCGTTGATGGGAAACAGTTTCAAGCCCCAGTCCCAATCACGAGCGGGGTTCAACGGGTTACCCGCGCCTCTCGGCGCAGGGTAGGCA

Hepg

si:ch211-106a19.1 (si:ch211-106a19.1), transcript variant X2, misc_RNA

AGCAGTCCATTGCTGAGCACGAACCATCTGCGCTGGTAACCCTTGATGTAGTTAGTCCATTTAAAGAGCCAGCCTTTGTAGGTGTCCGAGCCCGGGGCTGGTGTCCCGCTCGCCGGCGTGGACGCGGCACTCTTGCCCTGCTCGCTCATCCTCCTCCTGCCCCGCTCCACTGCCGCAGACTGATGGTGTTGCTGCCGCCGCTGCTGCTGCTGTTTGCAGGATAAAC

Hepg

neuronal tyrosine-phosphorylated phosphoinositide-3-kinase adapter 1 (LOC101885998), transcript variant X1

AAAATCAGACAACAACAAATCCACGCCGGTCTGCTGACGATGACATTGCCCTCTTCATAGAGGAAAGACGGAGCTGACAGTATTTCCAAACGTGTCATGAAAGATTAGACGCAAAATGTACGTCGGGAGACCGAGCATCAGTGCGCTGACGGCTGACTGCTTTTTGTAAACAACATCCTCATCCTCATTCACCCTCCGCGCGGTGCTCGATGAGAAAGCGCTTCGCTATCC

Chromosome6：

Hepg

serine/arginine-rich splicing factor 3a (srsf3a), transcript variant X3, misc_RNA

ACATCACGAAAAAAGTTGTACCTCAAAACCACCAAATAAATCAAATTCAATGTCATTAAGCGGAGGAAAAGGCGTTAACTTACTAACAAGCTAGTTATGATGCAGTGAAGCTGAGTGCTGGTCAGGAGGGCGAAATTTGCGGGGAGATTTTGCCAGATGGTTGCTAGATGTTAACTGCTATACAGTAGGTGGCTGAAGATGTC

Chromosome8：

Hepg

v-ski avian sarcoma viral oncogene homolog a (skia), mRNA

CCGAGTCAAAGCCCCAATGGCAAGTCCTGTTCTCCAGCGATTTGTGACTGTGCACCACGAATTTATGGGTCGGGTACATGAGCCTGCAGTCCATGCATTGGATGCAGGGCGCGTTGGGACTTGTGTAGAGCTCCGGGACCAGCAGACCCTTACACTTCCCGAAGCACTCGTGGTAGATCTTGAAGCTCTTCTCGGTGAGTTCCAGCTCAATAGAGCCGTACAGCTCCTTCTTGGAGTGCGGCGGGTACG

Homo sapiens SKI proto-oncogene (SKI), transcript variant X1, mRNA

Chromosome9：

293

nebulin (neb), transcript variant X23, mRNA

ACTCACATCACTCTGGAGATCATAGACTTTTCTGGCCTGAACGACGTCATTGGAATCAGGTAGGCAAGTCCAGTTGTGCAGGTAGGTCCTGTAATTGGCATCATTAACCTGGATCTGACACTTCTTTGCCAACTCAAGAGACAGAAGGTCCACTGGGAGGTGGAACTTGGTCTTGGACTTGTTGAAGTCTTTCTTGTAGAGAGCATCACTAGCAATCTTAGCTGAGTTCATGGCCA

 Homo sapiens nebulin (NEB), transcript variant X36, mRNA

293

sorting nexin 4 (snx4), mRNA

CATGCGTAATGATCTTTATCATCTGGCACGGAGGACACCAAGAGTCAGATTTCCTATGAAAAGTAAAGTAACTGTTCTTATTCACATGCAAACTAAATTGGTCCTGACGCCCATACAGCAAACAAAGTGCATAAATTCATCATAGTTTGCTGCTCTGTCACTTTTCTGCACGTTCAGCCATAAAAGCATAAGCTGTTTATGACCACCGGCCCCCTCCC

Hepg

myosin, light chain kinase a (mylka), transcript variant X3, mRNA

ATCTGCCGTTGGAGGTTCCCTCTGAAGTCCATCTGCTCAGCCGTGATCTCCTTCAGATCTTCCTCTGAGACGCTCTTAGTGGTGACCTTGCGGCCGAGCACGGTTCTGAAGTCGATCTGTTCGGCCTCCTGCTGTCGGATCTGATCTTCCCGATGCTCCTTCGTTTCTACACGCCGTTTGAGAAGCCCACGCGTGGAGCCCCCCTCCTCTTCTTGCTGATCTGTCCCACTGCTATG

Homo sapiens myosin light chain kinase (MYLK), transcript variant X9, mRNA

Hepg

speckle-type POZ protein-like a (spopla), mRNA

TTAACGTTTATTTGATGTCTAGTCGCACAACTAAACATTAGGAATATAATCTAGTATTTCCTGTGTGGATGAAACTCCTGGCGTATTCCTAGAGAGCTCCATATCAAAACAGCGAGACTTTGGGAACTAGTGCAACGAGGAAGAGGCCTGGTCTCACGGGACAAGGCAGGCACAGAGTAGAACTGAGAGAATTCAAAATACTCCACACAAGAGTCATGGCAGATAGTTTAAGGCTGTTTGAA

Hepg

interleukin 1 receptor accessory protein-like 1a (il1rapl1a), transcript variant X12, mRNA

TCATATGGTATGTATTCAAGCGGAGTTATTTGAGACTCTATACTAGGATGCGAGTATACAGGCCCGTGTTCCTCTGTGAAACATGCCAGCATGTATCTGAGTTACACAGCAGCTAGAGGAGAAAAAGACAAGCATGTAACCAGTGAGTTCAGATGTCACACTGAGAGCTAACAGATTAGAAACGTGAAAGAAGGTGAAAAGGT

Hepg

fibroblast growth factor 14 (fgf14), transcript variant X4, mRNA

GCATTCCATATCAGTTAGGGCATTCCATGGTGAAAGTTTAGCTGGTTTATCCAGTTGCATCCTCCTACGTGCACAGCTGACCACTCTTCCGGAGCCAGTCCGTCGTCGAATTAGATCGGAACAGTCAGAGGAGTGAAGAGAGGCGGAGGGAATGGGGAGGAGAGAGCAGAGGATGCCTGGATCTGATCCACAAACATGGAGAGTCTGACAAAAGGAGCACGGCTCTCCGGGGTTGAGCATGGCCG

Hepg

DEAD (Asp-Glu-Ala-Asp) box helicase 3a (ddx3a), transcript variant X15, mRNA

TTCTAGATTCTACTTTTGCTGCTAGTTTTGTGTAATTTATTAAGCATTTTTTGACAAATATTTATTTTTATAAGCCTCAAAGTGATTCTTTGAAAGTTTCAAGAAACTTGACCAAAAGACAATACCTAAACACTGGCACTTGAATGTTGAATGTCACCTGTATGCGTGAAAAATTTCTATATTTCGGGGTAGTGTGAGCTTTTTAATGTTTCAGACGTAATG

Homo sapiens DEAD-box helicase 3, X-linked (DDX3X), transcript variant X3, mRNA

Hepg

POU class 2 homeobox 1b (pou2f1b), transcript variant X9, mRNA

TCACGCTTTAAGAGGTCTTTATTTCGGAGGTCTTTGTAATTGCCTCTATTTCAAGATATACCAAAAATCTTTGTTATACCTGTTGCTGTATATTGCTACTGTAGCGTAGTGTAAGCCTAAATATTTATGAGCGAAGGCTCTCGGTAGACCACAGGAAATATAATGCAATATTTTGATCTTTTCCAAAGACGAATACATATTTTTCTTTCTTTTCTTTAGTTTTTTTTTATTATTATTTTAAGATGGCTGTCCCTCC

Hepg

low density lipoprotein receptor-related protein 1Bb (lrp1bb), transcript variant X4, mRNA

CGAAGTTTTGTGTATCCATTTAACCAGAGCCATGGACACAAAGTACATGCTCGCATCGAGGAAAATGCCCGCATTATTGGGATGGATGCTCTGTTTCACCAACAAAAGTTTATCTGGGCTACTCAATTTAATCCCGGTGGACTTTTTTACAAAGACATCCTAAACAGGAGTCAAACTAAAACAAATGTTGGCATCATAGTAAGTATCCTGAGTCTTGTGTGAAGCATA

Chromosome12：

Hepg

si:dkey-7e14.3 (si:dkey-7e14.3), mRNA

CCGCGTACAGAAGGTAACGAGAAAGGAAAAGGTAATTACTGGACGTTTGCCACAGGCTGTGAGTCCATGCTGGACCTCTTTGAAAATGGCAACTTTCGTCGCCGCCGCCGCAGACGCAACTTAAAAATGGGCCTGAAGGAACCAACTGAAGCCTTTATATCCATGGACGCTCATCAGGCCGTCGCAGTGAGATCCTCAGATTCAGATTCTTTCATGAACAC

Chromosome13：

Hepg

uncharacterized LOC110440175 (LOC110440175), ncRNA

ACCTGCAACTGCTACATTTCTGACACGGCGCATCCTGAATGAGTACAATTTACAGTAAGGAGATGAAAAATGTCATCTACAGGCATCTGACAACACGTCCAGCACTGATGATCAGACTCCATTTGTAACCTGTTCCCTTTTGTTGATTTGCCGTGTGATAAATTGCATTTAGCAAAGCGGGACAGGCTAAACGGTCCTGACAACTTTTATTGATTTGC

Chromosome15：

Hepg

nuclear receptor interacting protein 1a (nrip1a), transcript variant X1

ATTGCAGAGTGCCGGGGATGCCTTGAATAAACAATTCAGTCAAGCGACTGACTGCAGGACTCAAGAGCACTAGAAAACTGCACATGTGGGGAATAGACGCTGTCAGCCCCAGCCACTTTCCGTTTTTTTTAAAAACATCTGAATTGTTCGTAAAGAGAAATGTTATACGAGCCTCTCCGACTCTTCTCTTAAATGCAGCATATTTTTCAACGGGATTCTCTTCAGTCACCACCCGAACAGGTAAGGCTGGATGTGAAAATAGGATTTATTTATTTGTCTGTTGTAACAATG

Chromosome16：

Hepg

dual-specificity tyrosine-(Y)-phosphorylation regulated kinase 1B (dyrk1b), transcript variant X4

CACTTACCAGAGCTGGAGACAGAGCTCGTGGTGGAGGTGGAGTGGCTGTGGTCCATGGCAGGGCTGGTGGAGGTGGAGCTGCTGGTGTTGGTGCCCTCGTCTGTCGTCTTCTTAAAGAAGTTGTGCTGCAGCGCGTAAAAGGGCGTGATGCGCGTCTTGGGATCATAGTCCAGCATACG

Hepg

si:dkeyp-97a10.3 (si:dkeyp-97a10.3)

AACAAGCATCTGTGAGCAGAAAGCTGAATAAATAGTTGCGAGAGGACAAACAGTGGCAGTCCATTATTTATCATTTAGTAGTTCAGCTCTGCCAGCCGTCCTAATTCATTTGGATTTCGTTCAAGTCTTCCTGTCTGTGGAAAAACACTGCATTGTGTTTTATTTTCAGAGGTTTGAGTGCTAGAAAAAAGCCCAGAGCTGCATTGTTCTGAGGGATG

Chromosome17：

Hepg

kelch like family member 28 (klhl28)

TCACCTCTCCAGTGTGGCGAACAGTCCTGCTTTCCCCCCCACAGCGAGCAGTACTTTGGGAGCGCAGCGGGGCCGTGTGGACAGGACGGTCTGGTAGGAGAGCCGGTGTTCAGGCATGAAGTGGTACTTGAGCGCCTCGTTGAGCAGGTGTTTGCAGGCGTGGTCGTCCCGGATCAGGTGATTGGCCTCGTAGAGTCGCGTCAGGAACTTGACGCTCAGCAACGGCAGCCGGAC

Homo sapiens kelch like family member 28 (KLHL28), transcript variant X4, mRNA

Hepg

somatostatin receptor 1a (sstr1a)

ATGACAACAGATAACAAAAACGAGTCTTTATGCTGGCTGCAGTGCCTATAGTGTTGTAGTTCTGGATGTGCATGTGCCATTCCTGTACGTGCTGTCCGATTCCAGATTGTCCGGTTGAAAGTCATCCACGCTGTATCCCCGACTCTTCAGGGCCGTGGCATAGTAATCTATGGGCTCCTCCGTGGCGTTCTCCCACCATCGGAGGCATAAAATCCTCTGGAAGGAGCGTCTGAAGT

Hepg

discs, large (Drosophila) homolog-associated protein 2a (dlgap2a), transcript variant X3

GCACTTCGGTGACCTTTCACTAAAGACCTCTAAGAGCAACAATGACGTGAAATGTTCAGCATGCGAGAGCATCGCTATGGCCCCCGAGGGCAAATTCATGAAGCGCAGCTCCTGGTCGACTCTGACTGTCAGCCAAGCTAAAGAAGCCTACCGTAAATCTTCCCTCAACCTGGAGAAACCGACCATGCCTCAAGATCTGAAGTCCTCCATGAGGCCTTGTC

DLG associated protein 2 (DLGAP2), transcript variant 3, mRNA

Chromosome18：

Hepg

piccolo presynaptic cytomatrix protein b (pclob), transcript variant X6

GATGAGGATCAGGATGAGTGGGATGTTCCTGTTCGGAGTAGGCGCAGGTCTCGTTCCAGTAGATATGCTGATGGAGAGAAGGGTAAAAGCTCTAAGGTGTCAAGTATTGCGATTCAGACTGTAGCAGAGATTTCAGTTCAGACAGAGCATTCGGGAACCATTAGAAGATCTCCTGTTAGGGCTCAAGTGGACACAAAGGTAGATTTACAGAGAGAGGGCCAAACAGAA

Homo sapiens piccolo presynaptic cytomatrix protein (PCLO), transcript variant X1, misc_RNA

Chromosome19：

Hepg

bloodthirsty-related gene family, member 24 (btr24)

TCCCAGACAAACCACAGAGATTTGATTACTGTGTCTGTGTCCTGGGAAAGGAGGGATTCTCCTCAGGGAGATTTTATTTTGAGGTGCAGGTGAAGGGAAAGACTGACTGGGATTTAGGAGTGGTCAGAGAATCCATTAACAGGAAGGGACAGATCACAGCGAGTGCCAGTAAAGGATTCTGGACTGTGGTTCTGAGGAATGGGATTGAATATAAAGCCTGTGCTA

Hepg

si:ch211-233f16.1 (si:ch211-233f16.1), transcript variant X4

TGGCCTCTCCCATGTGGTAATATATAGCACAAAAATATAGGTGGTGAACGCTCCTCATATATATGCAAGTAGATTTACAGATCCCTTTCCCCTAAAAAAATACCCAGTCAACTGTGAAGAAGAAGAAGAAGCACTCACAGAGTGTGAATAAAGTAGCCGAAGGTTTTCTCCATGCCTTCCGTCATGAAGAAGCTTTTCAGAGCGAGTCTTTCAACAAAAAAAAAACCCCAACAAAAACC

Chromosome20：

293

eph receptor A7 (epha7), transcript variant X1

ATCAAACCAGAAAAGTTCAAAACGGCACAAAGCAGAAAAGGAGGAAAAAAAATTGACAGATTTGTCATTGACCCGTACGGTGGCCTCAATGTGGGATTGATTGAATCCAGGCGTTAATCTCAAATTGGCTCAAATTGCTGTCTCTACAGGGGCCCCGGAGAGGGTAAGGGCCTGTGAGGGGCCCTCGGGAGGCCGGAGTGTGAGATCAGAGCGCGGGGAGGCTGATCCGAGTGACTGCAGCCACTGCTGAAGCTC

Hepg

synaptosomal-associated protein, 25a (snap25a), transcript variant X1

TGCCACCATTTCAGTCGATATATATATTTACAGACTGTGACTTTTGTTGTAAAATAATGCAAAGTGACTTAAAGAAGCCGTGCGAAGGGATGGAAACAAAGCTGGAGTGGTGGTCCTGCAAAAACGACTTGTTTTCTAAAAATGAATATGCAGTTTGGTGTAGAAAATGGTGTTCTGAAAGGTGGGGACTGGTTGTTTACTGTATCATAGGTTTTGGATGCTCTGAAAAAAAAAAGAAATAAATCATTTGCA

Chromosome21：

293

crumbs family member 2a (crb2a), mRNA

TTCAATAACACTATTTTCAGTTGCTCGTTTTTGAGAGGGTTTTGTTGCTGGGAGAAAGTAAACAAATTTTTTTCCACCTCTTTACAGTATGACACTTTGTCAGAGCTGTTAAGGCATATAATGCCATGAGGTAAATCATCACCAAGTCCAATGTCATCTGGAAGAGTTTCTGGTTAACATCCACAGTCATATAATGACAGTTAATTCATATTAAATTCATTCCCATTTTCCAA

Chromosome22：

293

myosin heavy chain, fast skeletal muscle-like (LOC108181090), misc_RNA

ATTTCTGCTCCTGATCTCAGAGTCCAGAGTGCTCTGCATGGAATCCATCACTCTTTGGCTGTTCCTCTTGATCTGTTCTATCTCTTCATCCTTCTCAGCAAGCTTCCTGTCAATCTCACTCTTCACCTGGTTCAGCTCCAGCTGCACACGCAGAATCTTGGACTCCTCATGCTCTAGGGTGCCCTACAGGAGAAACAGACACAATCTAATTTCATTTT

Homo sapiens myosin heavy chain 1 (MYH1), transcript variant X1, mRNA

Chromosome24：

293

component of oligomeric golgi complex 7

GACCCCATAATGACACACTTTCAGTAAGGAAGAAATTCCGGCACACGTTCTTTACCGCATGATTAAACCCAAGTCTATAATATTATTTCCCTTTAAAGGGATAGTTCACACACAAATTGAATTCAGCTATCATTTACTTGCTCCAAACCTGTTTGAGTTTATCCAGTTGAGCACAAAAGATATTTTGAAGAATGCTTGAAACCTGTAACCATCGGCT
