## Supplementary material for "Zebrafish DANA retroposon can form large zebrafish sequence in human Hepg2 and 293T cell lines": table 3

Table 3. multi-copy fragments occurred in zebrafish genome

| Chromosome | 293 | Hepg2 |
| --- | --- | --- |
| 1 | 1 | 1 |
| 2 | 0 | 2 |
| 3 | 0 | 2 |
| 4 | 0 | 0 |
| 5 | 0 | 0 |
| 6 | 0 | 3 |
| 7 | 0 | 0 |
| 8 | 0 | 2 |
| 9 | 3 | 0 |
| 10 | 0 | 0 |
| 11 | 2 | 0 |
| 12 | 2 | 2 |
| 13 | 2 | 2 |
| 14 | 1 | 0 |
| 15 | 2 | 0 |
| 16 | 1 | 3 |
| 17 | 0 | 0 |
| 18 | 2 | 0 |
| 19 | 2 | 0 |
| 20 | 1 | 1 |
| 21 | 1 | 0 |
| 22 | 1 | 0 |
| 23 | 1 | 0 |
| 24 | 0 | 2 |
| 25 | 0 | 0 |

Chromosome1：

293 AATTAATCTGTTTGAGTAAACAAAGCAATTTGAGCACAGTAAAACCCAATAAATGAAGAGAACTCAAACCAACTGAGTTCTGTAAAACCCAATAAGTTAAGGCAACTCAAAATGTTTACGGAAACTGATCGCTACAAACTATTTGAGTTAAAAACCTAATTTATATGAGTACTGTAAACTTACTCCATTTAAGTTAAGGTAATGAGTAGTTTAATTCACTCATCACCTTCA

Hepg TTTCTTTTCGGCTTAGTCCCTTTATTAATTCGGGGTCGCCACAGCAGAATGAACCGCCAACTTATCCAGCACATGTTTAGCGCAGCGGATGCCCTTCCAGCTGCAACCCATCTTTGGGCAACACCCATACATACTCATTCACACTCATATATTATGAACAATTTAGCCTACCCAGTTCACCT

Chromosome2：

Hepg GAGTTCTGTGGATCTCAGAACTCCCCGGCTGCAGTAGCTAAGGGAACACGTGAGAAAAAGGGGGCTAGGCAAGTGCTTGATTGTTATGCATGATGTAAATCTTTGGATGGTGGGAGGAAACTGGAGAACACAGGGAAAACCCACACAGACACAGGGAGAATAAACAAACTCCTCACAGAACCACCCACCGGCCTGGCGGAACCAGGGTCGACAGTGTTCTTGATGTGA

Hepg CCATTTTGTGGATAGACAACGTCAACCGTCTTTGATTAATGGACACCGACTCTGTGTCTCTCAGAAATTATAAATATTTTGAAATAGGCAATATCCTTATAAATAAACTGCATAGTTGCAATCTAAACACCTACATCCTCGCCTAAAAAAAGCCCTAAAAGTACATTATGTTGCCCATCAGCAGGAATATTTGTCAAACTGTGGAGTGTCCTCTGCTATATGTGGGCGGAGTA

Chromosome3：

Hepg ATCTTCGGGCCTTAAGACGGCACTGCATCCCAAACAGGAATGCTACTATAATGTTAATCACAATATGGGCTCAGGACTACTTCCAGAAAACATTGTCGGTGAACACAATCGACCGTGCCATTCGCCGTTGTCAGCTAAAAGTCTATAGGTCAAAAAAGAAGCCATATTTAAACATGATCCAGAAGCGCAGGCGTTTTCTCTGGGCCAAGGCTCATTTAAAATGGACTGTGGGAAAGTGGAAAACTGTTCTGTGGTCAGACGAATCAAAATTTGAAGGTTTTTTTTTTTTTTTGCAAAACTGGGATGCCATGTCATCCGGACTAAAGAGGAAAGGGACAACCCAAGTTGTTATCAGCGCTCAGTTCAGAAGCCCGCATCGCTGATGGTATGGGGTTGCATGAGTGCATGAGGCATGGGCAGCTTACACATCTGGAAAGGCACCATCAATGCTGAAAGGTATATCCAAGTTCTAGAATAACATATGCTCCCATCCAGACGTCGTCTCTTTCAGGG

Hepg TATCATTTAAAGGTCCAGTCTTAAACTTTTTTTATTCCAGAATTTTAAGATAATCAGTTTTTAAAAAAGTAGCATTTTCCAGTTATTACAAACAAAACCCCATGAAGTTTTTTTTTCTTTACAAATACACACTATGATGTTTTATACTTCCAAAATAAATTTTGTCTACATCCACCTTTCCCTGAGCAAGTTATAGCA

Chromosome6：

Hepg

TTGAGGATTTGAAGAGGAGCTGTGTCCGTCCTTCTGTGCTTCTTCGGAAAACGAAGATAGGGGTGTACATCTTAGCTAGGTTATTTATAGTGGGTTTGGATCTTTCCGATTGGCTAGCTAATGAGCATAGATGAGTGGCCGGCTGCAGTCAAACATATCACGTGCTCCTCTCGAAATTAGTTTATGAAACTTCATTAAATGTTGCTCTCGATCCATCATGAAACTTGCA

Hepg AAGAGCTAATCTGTCCCACCCTCTCTCCATGTTTCAGTTGAGATTACGTCAAACATAGAATTAAAATGCTAAATTGAATCATCCAACTATGATGGAAAGACATTTACTGAAATAAATTCCAACATGCGCATCAAAGAAAGTCATGTAATTTTGTTTTAACAGATTATGTGATAACTAGGAGTGATGGTACAGCATTAGACAAACAAGTGGACCGACCACAATGTAGAGCA

Hepg ACATGAAAGCATGGACCCATTCTGCCTTGTATTAACTATTCAGGCTGCTGGTGGTGGTGTAATGGTGTGGGGGTATTTTCTTGGCACATATTGGGCCCATTAGTATCAATTGAGCATTGTGTCAACACAACAGCCTACCTGAGTATTGTTGCTGACCATGTCCATCCCTTTATGACCACAGTGTACCCATCTTCTGATGGCTACTTCCAGCAGGATAACGCACCATGTCATAAAGCTT

Chromosome8：

Hepg CAGCCCATCACATTACAAACCACAGGAGAGGGCAAATTCTTCAGCAGGATAGCGCTCCTTCTCATACATTAACCTCCTCATCAAAGTTCTTGAAAGCAAAGAAGGTCAAGGTGTTCAAGATTGTCCAGCCCAGTCACCAGGTATGAAGATTATTGAGCATGTCTGGGGTAAGAGGGAGAAGGCATTGAAGATGAATTCAAAGAATCTTGATGATCTCTGGGAGTCC

Hepg TTCGGTATTTACCTTCTTTTTTGCTGTTTTGCTCGCGCGAACAAATATATTTTTGTCAGCCGTGCTGCTTCTCGATTAGATCACATTTGCATGAATATTGAGCCATCTTTATTGTCGAACCCTGCTTTTAAGAAAACTTCAATTTAAAAATGATTTCCAACCAGCCAAAGTGGCTAGTGGAGGTGTCTGTTTAATTCTAACCCACCAGAGCTGAAATCTACCCTCATT

Chromosome9：

293

TTTGGCTGGACTCCCAGCTTGGCTGGGCTGGTTAACCTGGTTTTAGCTGGTCATCTCCCAGCTTGACCAGCTTAGACCAGGCTGGAAATGGCTGGAAACCAGCCTGGAAGGGGCCAAAACTCATCTAAAACCAGGCTGGTCGACCAGCTAAAACCAGCCAACCAGCCTAGGCTGGTTTAAGCTGGATTTTTCAGCAGGGTAGTGCTAACTTA

293

CAGCGTGGCTGGCGAGAAAGGGTATAAAAGAAGAAAAACTAATGACATGGCCTCTTTGTTCACCTGATCTGAACCCCATTGAGAACCTGTGGTCCATCATCAAATGTGAGATTTACAAAGAGGGAAAACAGTACACCTCTCTGAACAGTGTCTGGGAGGCTGTGGTTGCTGCTGCACGTAATGTTGATGGTGAACAGATCAGAACACTGACAGAATCCATGG

293

CTGAGAATAGGTGTTTATGTTCCTCTGCGTCGAGTTTTTTCGCTGGTGTTTTGTTTTTTCGGAACGCTTCCTTAATGAACAAGTGGTTCAAACTCGCTCATTTTGAGGCAGGAACCGGCAGGAATGCAACAACTTTAATCGTAATGTAAACACAAACCAACAGTCTCCATCCGGAGCTCCTTGTAAGTAGTCGCTCCATCGGGCTCACG

Chromosome11：

293

AGGACTGATTTTTCTAATACCCATAGACAAGAAAGACAAGTAGTGACTTCAGGTTGAAGGAAAGTAGTGGAGTAAAAGTACCAAAACGGCACTAATAATGTACTCAAGTGAAAGTAAAAGTAAATTTTTTTAAAACTACTTAGTAAATTACAATTCCTGAGAAAAACCCAATCACAGTAATTTGAGTATTTGTAATTAGTTACTTTACACCACCGGCTGTTGGCTAGAATTATGCATTAA

293

GATTGAGTTCTGGTGACTGTCGAGGCCATTTAAGTACAGTGAACTCATTGTCATGTTCAAGAAACCAGTCTGAGATGATTCACGCTTTATGACATGGGGCTTTATCCTGCTGGAAGTAGCCATGATAGATGAGTACACTGTGGTCATAAAGGGATGAACATGGTCAGCAACAATACTCAGGTAGGCTGTGTTGTTGACACAATGCTCAATCGGTATTAATACACCCAAAGTG

Chromosome12：

293

ACTTGACAGGCTCAGAAAAGTCAAAAATAGTGAGATATCTTGCAGAGGGATGCAGTACTCTTCAAATTGCAAAGCTTCTGAAGCGTGATCATCGAACAATCAAGCGTTTCATTTAAAATAGTCAACAGGGTCACAAGAATCGTGTGGAAAAACCAAGGCGCAAAATAACTGCCCATGAACTGAGAAAAGTCAAGCGTGCAGCTGCCAAGATGCCACTTGCCACCAGTTTGGCCATA

293

GGCCATGTCTCTGAGTATTGCACATCTTGTGCTTTTGGGCACTCCAGGGATGTTGCAGCTCTGAAATATGGCCAAACTGGTGGCAAGTGGCATCTTGGCAGCTGCACGCTTGACTTTTCTCAGTTCATGGGCAGTTATTTTGCGCCTTGGTTTTTCCACACGCTTTTTGCGACCCTGTTGACTATTTTGAATGAAACGCTTGATTGTTCGATGATCACGCTTCAGAAGCTTTGCA

Hepg GGACTTTCTTGGTCGCCTGAAGCCTTCTTTACAAGAATTAAACCTCTTTCCTTGAAGTTCTTGATGATTCTATAAATTGTTGATTCAGGTGCAATCTTAGTAGCCACAATATTCTTGCCTTTGAAGCCATTTTTATGCAACGCAATGATGGCTGCACGCGTTTCTTTGCAAGTCACATGGTTAACAATGGAAGAACAATGATTTCAAGCATCACCCTCCTTTTAAC

Hepg TACCTTGGCCATGTCTCTGAGTATTGCACATCTTGTGCTTTTGGGCACTCCAGGGATGTTGCAGCTCTGAAATATGGCCAAACTGGTGGCAAGTGGCATCTTGGCAGCTGCACGCTTGACTTTTCTCAGTTCATGGGCAGTTATTTTGCGCCTTGGTTTTTCCACACGCTTTTTGCGACCCTGTTGACTATTTTGAATGAAACGCTTGATTGTTCGATGATCACGCTTCAG

Chromosome13：

293

AGTAATAGTCACCTGCACACACAGATATCCCCCTAAAATAGCTAAAACTAAACTAAAAACTACTTCCAAAAATATTCAGCTTTGATATTAATGAGTTTTTTGAGTTCATTGAGAACATGGTTGTTGTTTAATAATAAAATGATTCCTCAAAACTACAACTTGCCTAATAATTCTACACTCCCTGTATATGCAATATGCCTGTTGCAAAGCGCTGCAATAACCTGTGTTACATGT

293

ATATTTGTGAATGTGGAGATGTTATATTGGTTTCACTGATAAAAATAAATAATTGAAATGGGTATATTTTTGTTAAGTTGCCTAATAATTATGCACAGTAATAGTCACCTGCACACACAGATATCCCCCTAAAATAGCTAAAACTAAAAACTACTTCCAAAAATATTCAGCTTTGATATTAATGAGTTTTTTGGGTTCATTGAGAAC

Hepg

AGGGTATAAAAGAAGAAAAACTAATGACATGGCCTCCTTGTTCACCTGATCTGAACCCCATTGAGAACCTGTGGTCCGTCATCAAATGTGAGATTTACAAAGAGGGAAAACAGTACACCTCTCTGAACAGTGTCTGGGAGGCTGTGGTTGCTGCTGCACGCAATGTTGATGGTGAACAGATCAGAACACTGACAGAATCCATGGATGGCAGGCTTTTGAGTGTACTTGCAAAGAAAGGTG

Hepg

TTGCTAAAATGCTTATGATAAACTGCTGTTTGTTTATCTTCAAGCTCTGCTTCAGTTGCTTGACGCGTGCTCTGGTGAGCGCCAGCACACACACATATTATGTACATCTTGACATTAGAAAAGTATTCGAAAGTATTCCTATATATGTTTTCATTGAAGTTATTCACAATACTTATCTATCCACAGAATTTGTAAAGAGGTCAAATGTGTAGACATTAAGC

Chromosome14：

293

AAAATTTGTTTCAAACCGGAAGTACAAATTTGCTCAAAATAACGCAAAAACAACAAATATATGTGACCTAATAGTGTTTTTAGCAGTGTGGGACACATATACGACTGTCAACAGTTCAAAAAATGCGTTTTGGTGTTTTGTGACCCTTTAAAAGCAAAGGGCGGCCCCTAGTGGTTCGGTGGTATGGGTTGCGTATAGGGAGGAT

Chromosome15：

293

CAAAGATTTGTATGGCTGCCAATGGAACTGGTTCTCTTGTATTTATTGATGATGTGACTGCTGACAAAAGCAGCAGGATGAATTCTGAAGTGTTTCAGGCAATATTTCTGCTCATATTAAGCCAAATGCTTCAGAACTCATTGGACAGTCCTTCAGCGTGCAGATGGAGAATGACTCAAAGAAACTGCAAAAGCGACCAAAGAGTTTTTG

293

TGTCCTTCTGCAAAAAAAGTCTGTACTGCAGAAATTCTGTGTCTGTATTGTTTTTTTACATAAACTGTAAATTTTATAATGCAATTTTTTTCTACGTTTTGTATAGAATTTTTGCAGTAAACGAAAGAAAATGCATGCATTTACTTAACCCAACGAAACAAAACTGTAACGAGACTTCAAATATAAGGGCAGAGCTGACACATAAACTAATGCAGTTTCTGTAATGTCAGCTGATGCTAGTGT

Chromosome16：

293

GGTGGTGTAATGGTGTGGGGAAGATTTTCCACAGCCTACCTGAGTATTGTTGCTGACCATGTCCATCCCTGTATGACCACAGTGTACCCATCTTCTGATGGCTATTTCCAGCGGCATAACACACCATAAAGTCATAAAGTGCGAGTCATCTCAGACTGGTTTCTTGAACATGACAATGAGTTCACTGTACTCAAATGGCCTCCACAGTCTCCAGAGCTCAATCA

Hepg

ACGGTTCCAGTTGGTGATGATTGTGCTGTCTCTACGGATTTGGTAAGTGTGTGAACTGTTCTTTGGTTTATTCAGATGGCAAAGTTACTTAGTATCATCAGCAGTACTCTTTCAGTTTTTAAAGACATACATTAGTCAAAATGATTTTGTTACTAAGTTTACAGTATGTTAATATCACTACAATATTAGGAATTGAGAGATCTGGTATAAATGCTGCTGCTGTAAACACCTGTACACATG

hepg

AGAAGTGTGAGTGCGCTGAATCGGGCTCAAGCACGGTTCACTTGGCCGGCCCTGGCCCGGTTGGAAGAGGTGTGCCTGAGCGCGGTACACTTGGGCTTTGGCGCGGTACGCTTGTGTGTGAGCGCGAAACGCGCCAAAGCCCGAAACCGAAAGCGAGACGTGACTTTTAAGGGACTGTTTCATATGGATTTATTAATCATTCTTACTGTTCAGTGATCGCAAACT

Hepg

ATGTGAATCTCCCATTCCACCACATCCCAAAGATGCTCTATTGGATTGAGAACTGGTGACTGTGGAGGCCATTTGAGTACAGTGAACTCATTGTCATGTTCAAGAAACCAGTCTGAGATGATTGCTGGAAGTAGCCATCAGAAGATGGGTACACTGTGGTCATAAAGGGATGGACAT

Chromosome18：

293

ATCAGTCTTGAGATATTTCTTGGCCCAGTCTTGACGTTTCAGCTTGTGTGTCTTGTTCAGTGGTGGTCGTCTTTCAGCCTTTCTTACCTTGGCCATGTCTCTGAGTATTGCACACCTTGTGCTTTTGGCCACTCCAGTGATGTTGCAGCTCTGAAATATGGCCAAACTGGTGGCAAGTGGCATCTTGGCAGCTGCAC

293

TGTGAGGTATCAAACATTCCAGTGTCATTAAACGCAGTCACTGTCTTTCTCCCCTGCGTCTGTGTGTTTGTTTTGCCTCGGTGAAAATCAGCGCGTGCCCAAATGGAAACTCCCATTTTTATCCAAATCCTTCCCTCTTTCCCTCCCCGGACACTCCCACCTTAACAGAACTTGACACACCCACTTTCCTGACTTTTTCCAAACTAGAGGTGT

Chromosome19：

293

ATTCTTCTTTCGTTCTATTTCGTTCTTATTTAAGCCTGTTTGGTGATTTCAGGCATCATACAGGCGACATCCTTTATTTTCTCTTCAGTGACATGCAGTCTATAAATTGTCTCTGGGTCATTGCGTGTAAAGATTTTGAACATCTTCTTGGACATTTTTAATGCTTCCAAACAGTTTGCTGCATTTATAAAGCGCCTATGTCTTGAGGTTGTTCAGTATGATGCAATTTACTTTCT

293

CCACAATCCAAAGACATGCGGTACAGGTGAATTGGGTAGACTAAATTGTCCGTAGTGTATGAGTGTGTGTGTGTGAATATGTGTGTGGATGTTTCCCAGAGATGGGTTGCGGTTGGAAGGGCATCCGCTGTGTTAAAACTTGCTGGAAAAGTTGGCGGTTCATTCCCCTATGGTGACCCCGGATTAATATAGGCACTAAGCCAAAAAGAAAATTAATAGGGCTGCAACAACAAATCGATTAAA

Chromosome20：

293

AATAATGTTGATCCTGGACCATCATCATCATTTAAATACAGTGTAAATTCACTAATGTTGATCCTGGACCATCATCATTTAAATACAGTGTAAATTCACTAATGTTGATCCTGGACCATCATCATCATT

Hepg TTCAGCCTTTCTTACCTTGGCCATGTCAAGTATTGCACACCTTGTGCTTTTGGGCACTCCAGTGATGTTGCAGCTCTGAAATATGGCCAAACTGGTGGCAAGTGGCATCTTGGCAGTTGCACGCTTGACTTTTCTCAGTTCATGGGCAGTTATTTTGCGCCTTGGTTTTTCCACACGCTTCTTGCGACCCTGTTGACTATTTTGAATGAAACGCTTGATTGTTCGATGAT

Chromosome21：

293

CATTTTAAATCAAGTCCAATCAACTGAATTTACCACAGGTGAACTCCAATTAGGCTGCTAAAACATCTCAAGAATGATCAGTGGAAACAGAATGGACCTGAGCTCAATTTAGCGATTTACAGCAAAGGCTGAGAATACTGATGTACATGTGATTTTTCAGCTTTTTTATTTTGAATAAATTTGCAACAATTTAAAAAAATCTTTTTCACATTGTCATCAAGGGGTATTG

Chromosome22：

293

TGTTTTTGCGTTATTTCGAGCAAATTCATTCTTCCGGTTTGAAACGAATTTTTGAAGCTGCGTCACGCCGATTAGATACTGGCATGTATTCCAGCGTGTAGACAGGCTGTCTTTATCTGGGAGTACTTTGTGACGTCTCTGGGTGTGCTGCATTAATTCATGAGAAAGGCTGGGTTTAAACCAATCAGTGCGATCTATTGTGTATGCAGTGATGCAG

Chromosome23：

293

TTCTAAACATAATAGTTTTAATAACTGATTTCTTTTATCTTTGCAATGATGACAGTGAAAAATATTTTACTTGATATTTTTCAAGACACTTCTTTACAGCTTAAAGTGACATTTAAAGGCTTAACTAGGTTAATTAGGTTAACTAGGCAGGTTAGGGTAATTTATAGGCAAGTTATTGTATAACGATGGTTTGTTCTGTAGACTATCGGAAAATATATA

Chromosome24：

hepg

TTCAAGAAAAACATGATTTTCATGCAGGACAATGCTCCATCACACGCATCCAAGTACTCCACAGCGTGGCTGGCCAGAAAGGGTATAAAAGAAGAAAAACTAATGACATGGCCTACTTGTTCACCTGATTTGAACCCCATTGAGAACCTGTGGTCCATCATCAAATGTGAGATTTACAAAGAGGAAAAACAGTACACCTCTCTGAACAGTGTCTGGTAGGCTGTGGTTGC

hepg

TAGCCATATATTTTAAATAATGTGTCAAAAATAATGTCTTTGAATCTAAAATTTAAATATTTTTTAAATGTTCAAATATTTAGCAGTAAAATACAGCAGTATGGAAAATGTGCAGGTATTTTATTGCTAAACAAAAGGTGGCTGACTTAAAAGTGATGGAAAACATTTTTTTCGTTGTGTTGTACCCATGTAAATATCTTTTTCCACAAAGTTTTAA
